# Cell-scale autonomous CMOS motes for intracellular bioelectronics

**DOI:** 10.64898/2026.05.12.724193

**Authors:** Girish Ramakrishnan, Filipe Cardoso, William Stoy, Victoria Andino-Pavlovsky, Joaquim Goes, Kenneth Shepard

**Author notes:** These authors contributed equally to this work (Girish Ramakrishnan, Filipe Cardoso).

## Abstract

Integrating autonomous electronics within single cells has remained beyond the reach of modern bioelectronics. Miniaturization at this scale could transform our ability to study and actuate biological processes at the cellular level, complementing existing molecular and fluorescent approaches. As these devices approach the sub-100-µm length scale, volumetric constraints demand fundamentally new approaches to power delivery and telemetry. Here, we report an optically powered 10-picoliter complementary metal oxide semiconductor (CMOS) mote that operates with a power density of 1 pW/pL, comparable to the metabolic rate of cellular systems. These fully CMOS motes can be manufactured at scale yielding 1000 motes from a 4-mm^2^ silicon die. Multiple motes can be simultaneously powered and interrogated within a single optical field of view using epifluorescence microscopy. We demonstrate intracellular implantation of these motes within the single-celled mixotrophic dinoflagellate *Noctiluca scintillans* with negligible cytoplasmic displacement, pushing the boundaries of active CMOS bioelectronics to the intracellular domain and establishing a next-generation of truly cell-scale bioelectronic interfaces

**Teaser:** A 10-pL autonomous CMOS mote with fluorescence-based backscatter communication enables cell-scale sensing.

## Introduction

There is growing interest in technologies that can enable precise sensing and actuation at the single-cell level. Techniques such as patch-clamp electrophysiology, quantitative phase and Raman imaging, and nanopipette-based electrochemical or ionic measurements capture dynamic changes in membrane potential, ion flux, and biochemical composition within individual cells (*1–3*). Complementary methods including single-cell omics (scRNA-seq, ATAC-seq, proteomics) (*4*, *5*) and live-cell reporters provide high-resolution single-cell views of transcriptional states and molecular signaling pathways. These and hybrid approaches that combine them bridge fast, real-time electrical and optical readouts with high-content molecular profiling, offering both temporal precision and mechanistic depth (*6*, *7*). Parallel advances in chemogenetic and optogenetic tools allow targeted manipulation of intracellular pathways (*8*), while nanoelectroporation and nanopipette delivery enable localized introduction of nucleic acids, proteins, or nanoparticles directly into single cells (*9*). Collectively, the ultimate goal of these technologies is closed-loop interrogation of intracellular dynamics, in which sensing and stimulation are integrated to control gene expression, metabolism, and signaling in real time within living cells.

Here we consider how intracellular bioelectronics, enabled by the rapid scaling of CMOS technology, could be an important new platform for closed-loop interrogation of cellular processes. To date, intracellular devices, have been exclusively passive, incorporating no active electronic devices or circuitry. This includes silicon-based passive microchips for intracellular pressure monitoring (*10*), bar coding (*11*), and passive radio frequency identification (RFID) tagging (*12–14*). Small active electronic chips have been pursued in other contexts (*15–19*), in which some of the challenges associated with wireless power delivery and communications, have begun to be addressed. However, these devices have not yet been reduced to deep-sub-microliter scales, compatible for intracellular delivery. At these dimensions, both power transfer and telemetry must be wireless and highly efficient.

Optical interfaces, in particular, are attractive for devices smaller than 100 μm. This is because typical optical wavelengths in the visible spectrum are orders of magnitude shorter than the wavelengths of alternative modalities such as radio-frequency electromagnetic and acoustic transduction allowing for aggressive miniaturization of the transducing element without substantial loss in transduction efficiency (*20*). Moreover, silicon, is intrinsically photo-active, allowing easy monolithic integration of optical sensing elements within standard CMOS processes.

In this article, we demonstrate a monolithic fully autonomous sub-10-pL CMOS electronic mote measuring 30 μm × 30 μm × 10 μm. This device functions as a temperature sensor over the temperature range of 20 ℃ to 30 ℃. Powered entirely by harvested light, the mote operates at a peak power dissipation of 10 pW. Power harvesting and telemetry are performed directly on the CMOS die, where the latter is accomplished with backscatter communication with a monolithically integrated optical fluorescence-based modulator. Systems of this size must be manipulated in liquid suspension; we demonstrate fabrication techniques that allow release of motes from CMOS bulk silicon, employing conventional lithography and plasma processing techniques, enabling scalable production of thousands of motes from a single CMOS die. We demonstrate that these motes are small enough to be embedded into the cytosol of the single-celled mixotrophic dinoflagellate *Noctiluca scintillans,* extending active CMOS bioelectronics into the intracellular domain.

By operating in conjunction with widefield epifluorescence microscopy, real time bio-geographical imaging is possible. The mote harvests energy directly from the microscope’s excitation light, while its modulating optical fluorescent signal, is observable in the same optical field of view. This means multiple motes in the microscope field of view can be interrogated concurrently, allowing simultaneous readout using conventional scientific CMOS imaging cameras.

## Results

### Mote design

An illustration of the mote and its application as an intracellular sensor are depicted in **Fig 1a**. The mote consists of an integrated circuit fabricated on a 65-nm bulk CMOS process and a monolithically integrated thin-film fluorescent quantum-confined Stark effect (QCSE) modulator that enables optical wireless telemetry. The mote interfaces with a widefield epifluorescence microscope using harvested light for power. Uni-directional telemetry is established using the readout camera by means of the modulated fluorescence uplink. The power delivery and the data telemetry are on spectrally separate channels. Blue light (∼480 nm) is harvested by a photodiode for energy and also excites the QCSE modulator, whose modulated fluorescence (540 nm) is used for communication. The QCSE is driven by a temperature-dependent relaxation oscillator, allowing the mote to act as a temperature sensor. The overall block diagram of the mote is shown in **Fig. 1b**, showing the photodiode for energy harvesting, the relaxation oscillator, and the QCSE modulator. Approximately half of the active silicon area is occupied by the photodiode with the other half occupied by the CMOS electronics. Two 12 μm × 12 μm metal pads laid out over the active electronics constitute the electrical interface to the mote (**Fig. 1c**).

**Figure 1.**
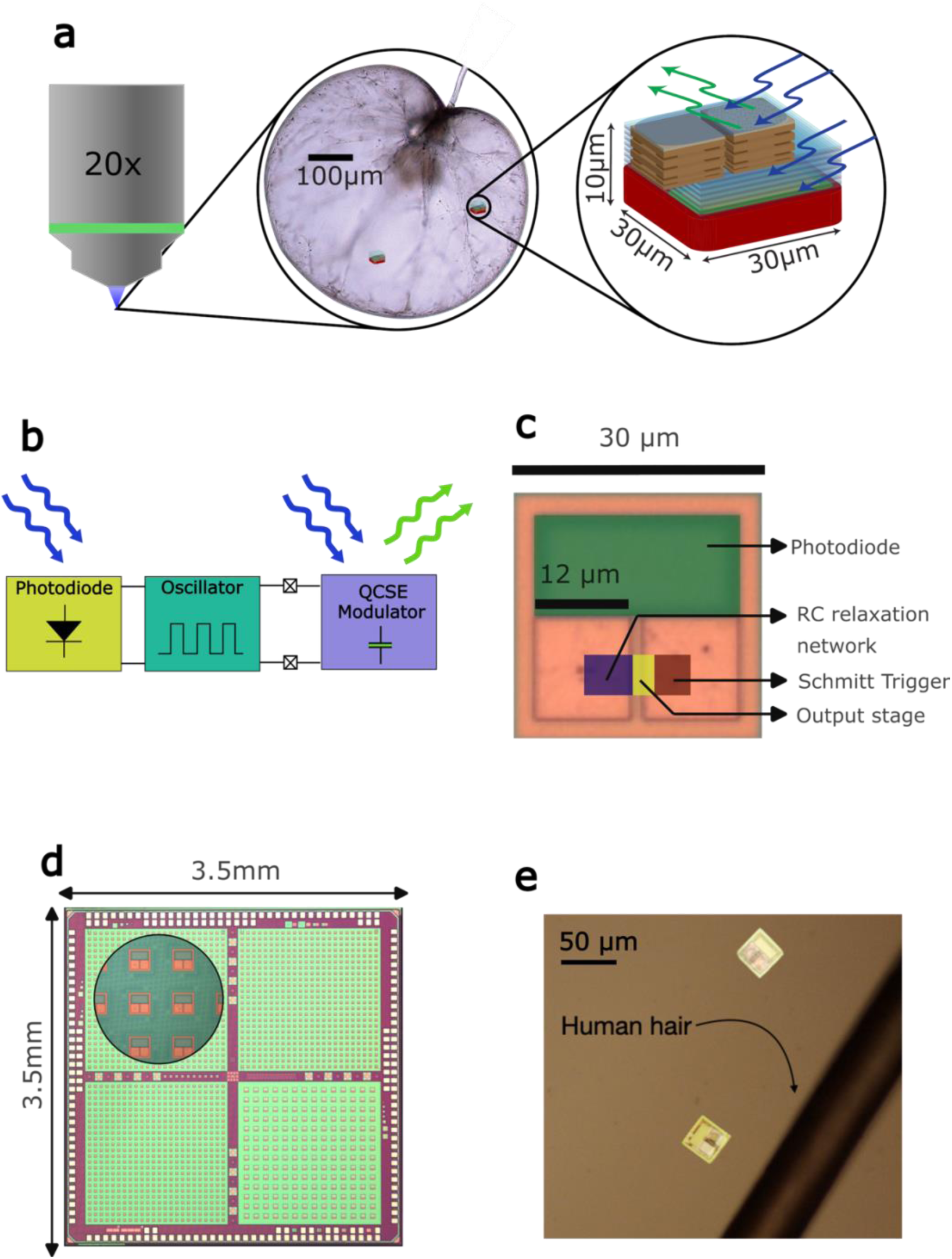
Autonomous micron-scale CMOS optical motes. **(a)** Illustration of an application for an autonomous intracellular sensing mote. The mote is powered under a widefield epi-fluorescent microscope and interrogated via fluorescence emission; **(b)** Block diagram of mote operation showing optical power delivery and fluorescent backscatter telemetry. **(c)** Micrograph of a single mote highlighting the active circuitry and the power harvesting photodiode. The oscillator voltage output is exposed via metallic pads on the chip surface. **(d)** Fabrication of motes at scale on a grid in a conventional bulk CMOS process. Monolithic integration and at-scale miniaturization enable manufacturing few thousands of motes on a die and millions of motes on a wafer; **(e)** Two motes adjacent to a strand of human hair.

All the circuitry is laid out within a separate deep n-well (**Fig. 1c**, **fig. S1a-c**), isolating the transistors from stray photo-generated carriers in the substrate. Metal layers are used to further shield the circuits from incident light on the top side. Incident light from the back side does not affect the active electronics because of the choice of short wavelength blue light for power delivery, which is absorbed close to the silicon backside surface before photons can access the active circuitry. The detailed layout of the motes on the die is shown in **fig. S1 and Fig. 1d**. Actuation of the modulator is accomplished with a relaxation oscillator whose time constant is determined by transistor gate leakage onto a capacitor. Two versions of the mote, each with a different type of relaxation oscillator, a gate-leakage-based relaxation oscillator (GLRO) and a gate-leakage-based three-stage ring oscillator (RO), were designed (*21*, *22*), as shown in **fig. S2**. The GLRO motes are designed to operate at 0.5Hz and the RO motes are designed to operate at 6 Hz. Additionally, the GLRO mote is designed in two size variants - a 30 μm × 30 μm variant and a 50 μm × 50 μm variant. The 50 μm variant uses the same circuit with a larger photodiode. Each die contains 1152 GLRO motes 30 μm × 30 μm in size, 196 GLRO motes 50 μm × 50 μm in size, and 576 RO motes 30 μm × 30 μm in size. All mote variants operate with a static power consumption of under 10 pW from the photo-induced supply voltage of approximately 0.4V on the photodiode. At 1 pW/pL the volumetric power density of these motes is comparable to the metabolic power density of similarly sized heterotrophic free-living protozoans in homeostatic steady state (*23*). The relative size of the motes compared to a strand of human hair is shown in **Fig. 1e**.

The parts of the mote are highlighted in **fig. S3**. The photodiode is implemented using two parallel p-n junctions, a shallow pwell-nwell junction and deep nwell-substrate junction that together implement a large photon capture cross-section using all the available volume in the CMOS process (**fig. S3b**) (*24*). The shallower junction absorbs the shorter wavelengths, and the deeper junction absorbs the longer wavelengths, conferring a broadband spectral response to the photodiode enabling efficient photon capture at multiple wavelengths. The interlayer dielectric stack of the backend of the CMOS process, however, does create a thin-film interference filter, causing a strong dependence of quantum efficiency on wavelength. The overall measured spectral response of the photodiode resulting from these effects is shown in **fig. S4**. To minimize optical loss due to thin-film reflections from the stack, the mote was operated using a light source at 470 nm.

### QCSE modulator design

The use of light emitters, such as light-emitting diodes, for data telemetry is precluded by both the power envelope and the volume requirements of this mote design. We instead employ a backscatter approach in which the energy for the link is not drawn from the mote’s circuitry. This is accomplished by an optical backscatter communication link which is implemented using a fluorescence modulator. The modulator is implemented as a quantum-dot layer contained within the dielectric of a parallel plate capacitor with a conductive transparent top plate (*25*) made from indium tin oxide (ITO) as illustrated in **Fig. 2a**. The quantum dots employed in this device are core-shell CdSe/ZnS quantum dots that emit at 540 nm embedded in a hafnium-dioxide high-κ dielectric in the capacitor. The voltage across the capacitor driven by the mote circuitry of amplitude ∼0.4 V, polarizes the quantum dots. This leads to a quenching of their fluorescence response by reducing the overlap integral between the electron and hole wave functions (**fig. S3d**), which has the effect of reducing the recombination likelihood of the electron-hole pair and causing a drop in the fluorescence response which is proportional to the voltage applied (*26*). Measurements from an externally actuated standalone QCSE modulator fabricated by thin-film processing is shown in **fig. S5**.

**Figure 2.**
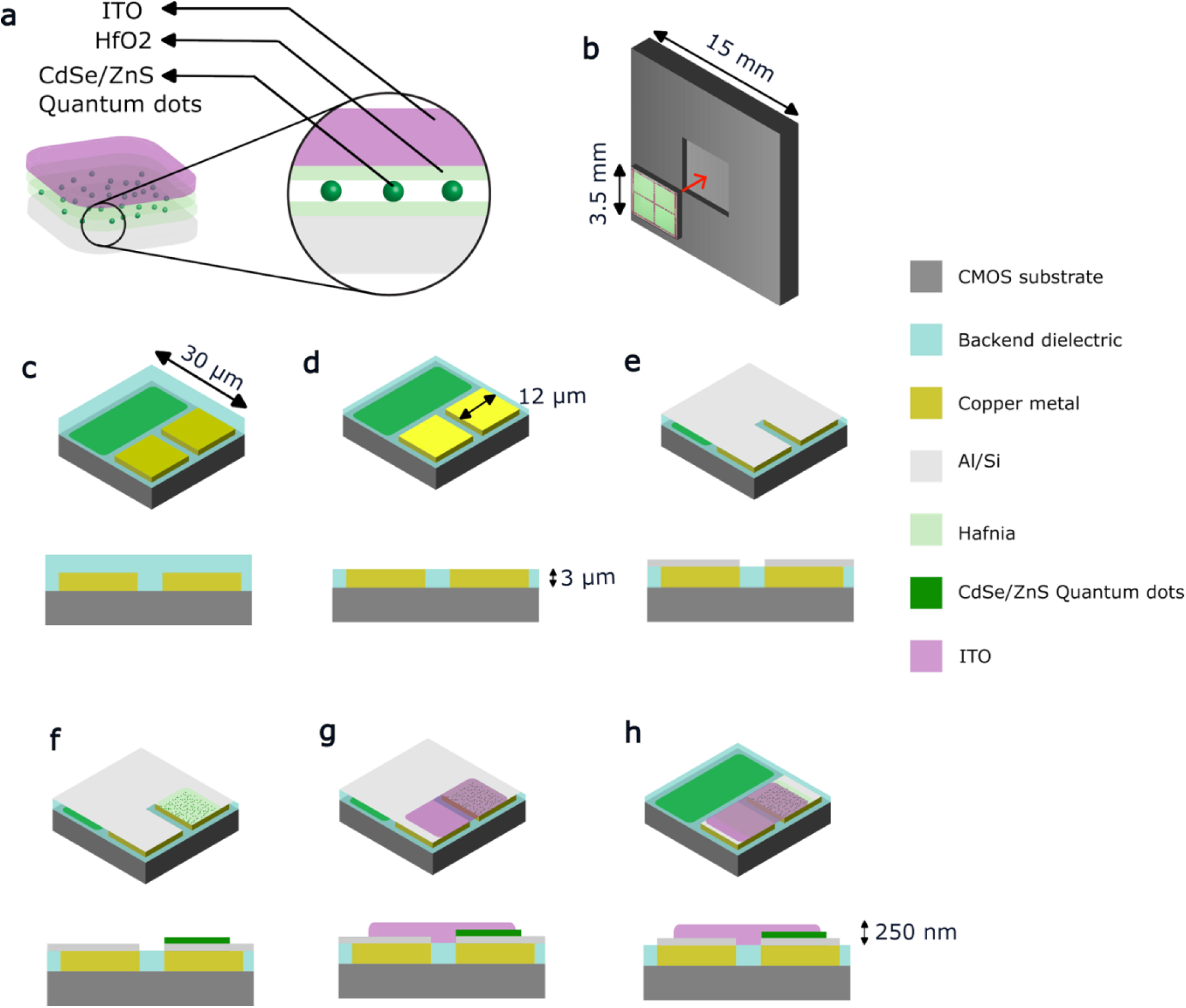
QCSE modulator fabrication. Isometric view and cross-section shown. (**a**) Cross-section and cartoon of the QCSE modulator structure. **(b)** The attachment of the CMOS die to a custom handle chip with a recessed pocket. **(c, d)** Removal of the top dielectric passivation layer to expose the copper pads. **(e)**. Deposition of a Cr/Al metallic interface layer. **(f)** Deposition and patterning of a Hafnia/QD/Hafnia layer on the device. **(g)** Deposition and patterning of the top ITO layer. **(h)** Final etch of Cr/Al interface layer to expose the photodiode and complete the modulator fabrication.

The process of fabricating the modulator by sandwiching the quantum dots within ALD hafnia films caused a significant loss of nearly 80% in the photoluminescence quantum yield (PLQY) of the quantum dots after the deposition of the top hafnium-dioxide film. This was likely because of chemical interactions between the precursors used in the ALD process and the unpassivated quantum dots leading to surface traps that activate non-radiative recombination pathways. Passivating the quantum dot films differently could recoup a significant amount of this loss which could allow for more power-efficient designs or designs with higher effective communication rates.

### Fabrication and handling of motes

As shown in **Fig. 1d**, these motes can be manufactured at scale on a traditional bulk CMOS process. Mote copies are laid out on a gridded fashion on a CMOS chip; in our case, we have close to 2000 motes implemented on a single 3.5 mm × 3.5 mm die. A 30-μm-wide metal-free kerf region between the motes allows for etching to singulate the motes as described below. After receiving the die from the foundry, a “post-processing” fabrication flow is performed to make the QCSE modulators (**Fig. 2**), followed by singulation and release the motes (**Fig. 3**). The whole micro-fabrication process is low-temperature (< 300° C) to ensure compatibility with the underlying CMOS substrate. An overview of the processing flow is given here (see **Methods**).

**Figure 3.**
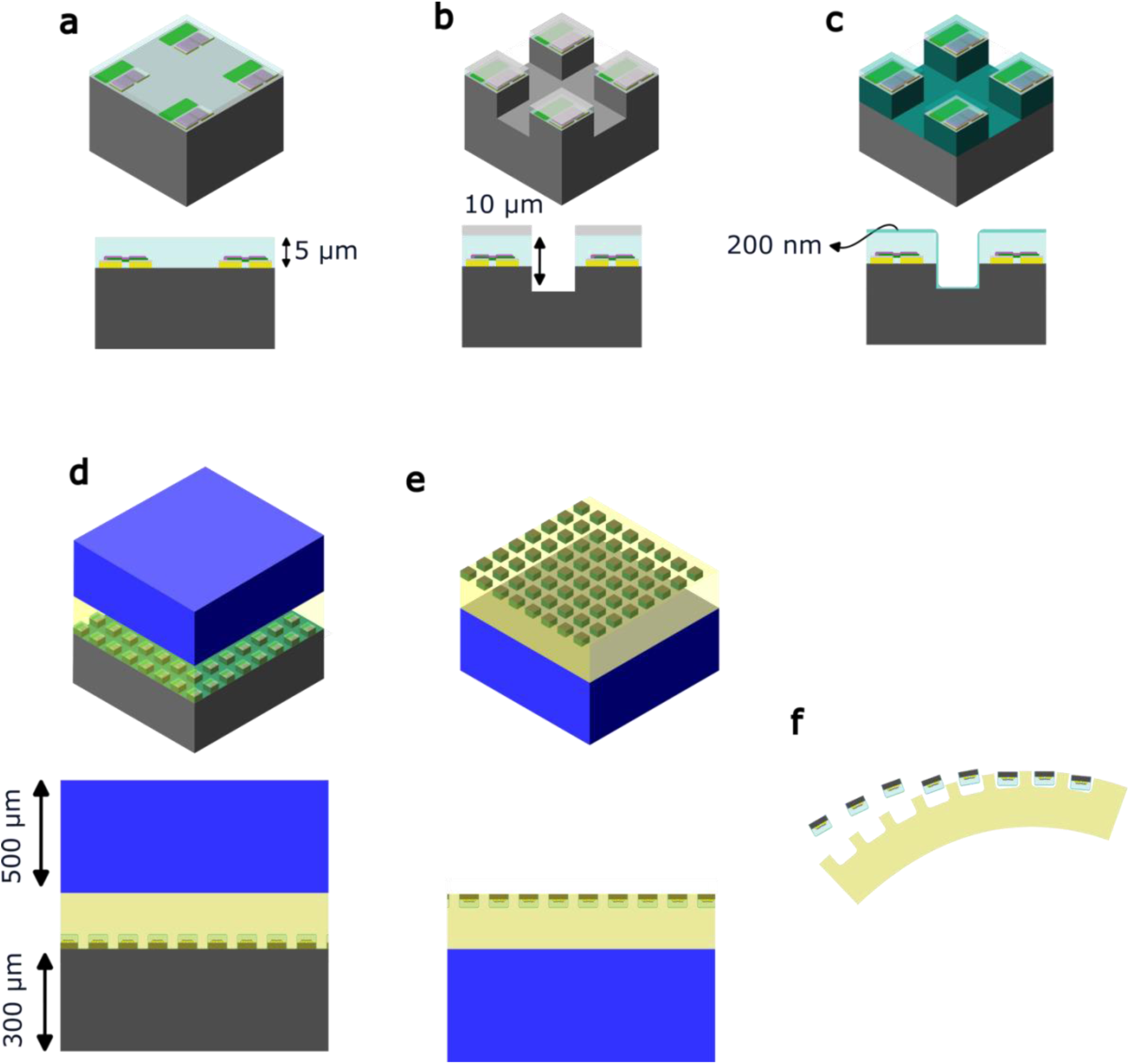
Release of the motes from the Silicon chip. Isometric view and cross-section shown (a, **b)** Silicon trench etch under a photoresist mask to define the mote regions. **(c)** Removal of etch mask and sidewall passivation using an SiO_2_ conformal barrier coating. **(d)**. CrystalBond support structure and handle wafer attachment. **(e)**. Backside silicon substrate etch to singulate the motes. **(f).** Release of motes from support structure into solution.

The modulator processing flow starts with mounting the die on a handle Si wafer into which a pocket is etched with the exact dimensions of the CMOS chip (3.5 mm × 3.5 mm × 0.250 mm). (see **Methods** and **Fig. 2b**). This helps with handling with the CMOS die easily and provides a flat work surface which is exceedingly important for defining the fine lithographic patterns that are necessary for manufacturing the modulator. The CMOS die from the foundry comes passivated with approximately 2-μm-thick oxide and nitride passivation (**Fig. 2c**) This passivation layer is removed to expose the top-level Cu interconnect of the CMOS die (**Fig. 2d**). The CMOS diffusion barrier layer is a natural etch stop for this process and guarantees a surface roughness of under 50 nm at the die surface, as measured with atomic force microscopy (AFM). To create the bottom contact of the QCSE modulator, we deposit 150 nm of Cr/Al and pattern it as shown in **Fig. 2e**. The dielectric stack of the modulator is formed by sandwiching a thin film of quantum dots between dielectric layers. First, a 10-nm-thick layer of HfO_2_ is deposited using atomic layer deposition (ALD) to form the bottom dielectric layer. This is followed by a layer of colloidal quantum dots which are spin-coated on the device to form the active optical layer. This is then capped with an additional 10-nm-thick HfO_2_ layer to complete the dielectric stack. The entire stack is then patterned to define the modulator device over a single metallic pad (**Fig. 2f**). A 100-nm-thick ITO film is sputtered and patterned to define the transparent top plate of the QCSE modulator. This top plate makes contact to one of the pads such that the parallel plate capacitor of the modulator is biased by the voltage between the two pads (**Fig. 2g**). Next, the aluminum film deposited in **Fig. 2e** is patterned as shown in **Fig. 2h** to expose the photodiode and the kerf region for downstream processing. Finally, we passivate the whole chip with a 1-μm-thick layer of SiO_2_ by plasma-enhanced chemical vapor deposition (PECVD) to encapsulate the motes and enhance their survivability in solution (**Fig 3a**).

The CMOS motes are finally singulated from the underlying silicon substrate by the process illustrated in **Fig. 3**. First, we define the mote regions with a stacked Photoresist/Cr etch mask. Trenches are defined by removing the backend dielectric material between the motes (**Fig 3b**). These trenches are 15 μm deep extending 10 μm into the Si substrate. The Cr hard mask is stripped, and the surface and the sidewalls of the trenches are passivated with a 0.2-μm-thick SiO_2_ layer to protect the implants post release (**Fig. 3c**). The motes are prepped for release by adhering the wafer face down to a glass handle with CrystalBond 509 (**Fig. 3d**). The CrystalBond forms a support structure to hold the motes after singulation. After bonding, the substrate is etched to meet the depth of the trenches such that the individual motes no longer share a common silicon substrate (**Fig. 3e**). Finally, the conformal SiO_2_ passivation is etched to completely singulate the motes. Finally, the motes are released from the glass handle using acetone into an Eppendorf tube (**Fig. 3f**). Since Si is denser than acetone, the motes form a pellet at the bottom of the tube (**fig S7a**) but can be easily resuspended with agitation. We centrifuge the suspension and replace the acetone solvent with an inert organic solvent, N-methyl-2-pyrrolidone (NMP). After three washes with NMP to remove residual organics, the motes are stored in the same solvent at 4 ℃. Some of the experiments conducted with the motes were performed up to a year after release and the motes continue to be functional even a year after release into NMP. For some of the experiments, we replace the NMP solution with an aqueous buffer solution or distilled water. More details of the release process are given in **Methods**.

### Optical and electrical characterization

An epifluorescent microscope was used to interrogate the motes under illumination. The motes were powered with widefield blue excitation light (470 nm) from the microscope (8 nW/μm^2^) and the chips were imaged with a green emission bandpass filter (535 ± 30nm). A CMOS camera with a 50-ms exposure time was used for data capture. The motes were first characterized collectively on the bulk silicon substrate before singulation by capturing time-series images, scanning across the chip with multiple motes in a single field of view. For example, the 576 GLRO motes on a single chip were captured by imaging 77 motes at a time in a single field of view (**Fig. 4b**) and scanning the microscope field of view across the chip. Five chips were scanned in total for a total of just under 3000 motes tested. The captured image data was pre-processed to remove the baseline fluorescence response and was de-noised using spatial averaging and temporal filtering. A Fourier transform was employed to identify oscillation frequency (*f_oscillation_*), allowing functional chips to be classified based on a signal-to-noise-ratio (SNR) threshold of 20 dB. The spatial distribution of *f_oscillation_*, observed on a full scan of the 576 GLRO motes on a chip is shown in **Fig. 4a** with non-operational motes in black. Across this chip, 422 motes were identified to be operational out of 576 motes, most operating between 0.25 and 0.75 Hz. Selected oscillatory responses are shown in **Fig. 4c** color coded against the relevant motes in **Fig. 4b**. Overall, across five tested chips, we were able to identify 1657 operational GLRO motes, translating to a process yield of 57% for the modulator fabrication. The histogram of the oscillation frequency over the 1657 operational motes is shown in **Fig. 4d** with 80% of all tested motes exhibiting oscillations between 0.25 Hz and 0.75 Hz. The solid black line in **Fig. 4d** represents the predicted distribution of *f_oscillation_* from a Monte Carlo circuit simulation of the mote circuit with the gate oxide thickness of the leakage device modelled as normally distributed with a standard deviation of 0.7Å. The exponential dependence of the tunneling current on oxide thickness (*27*) results in this significant variability in *f_oscillation_*. The RO motes were similarly tested across a single chip with an observed process yield of 27% (158 working / 576 imaged) with measured *f_oscillation_* between 2-8 Hz across the tested motes. The lower yield of the RO motes was likely due to non-uniformity in the modulator fabrication process across the die tested rather than a systematic difference between the two mote designs.

**Figure 4.**
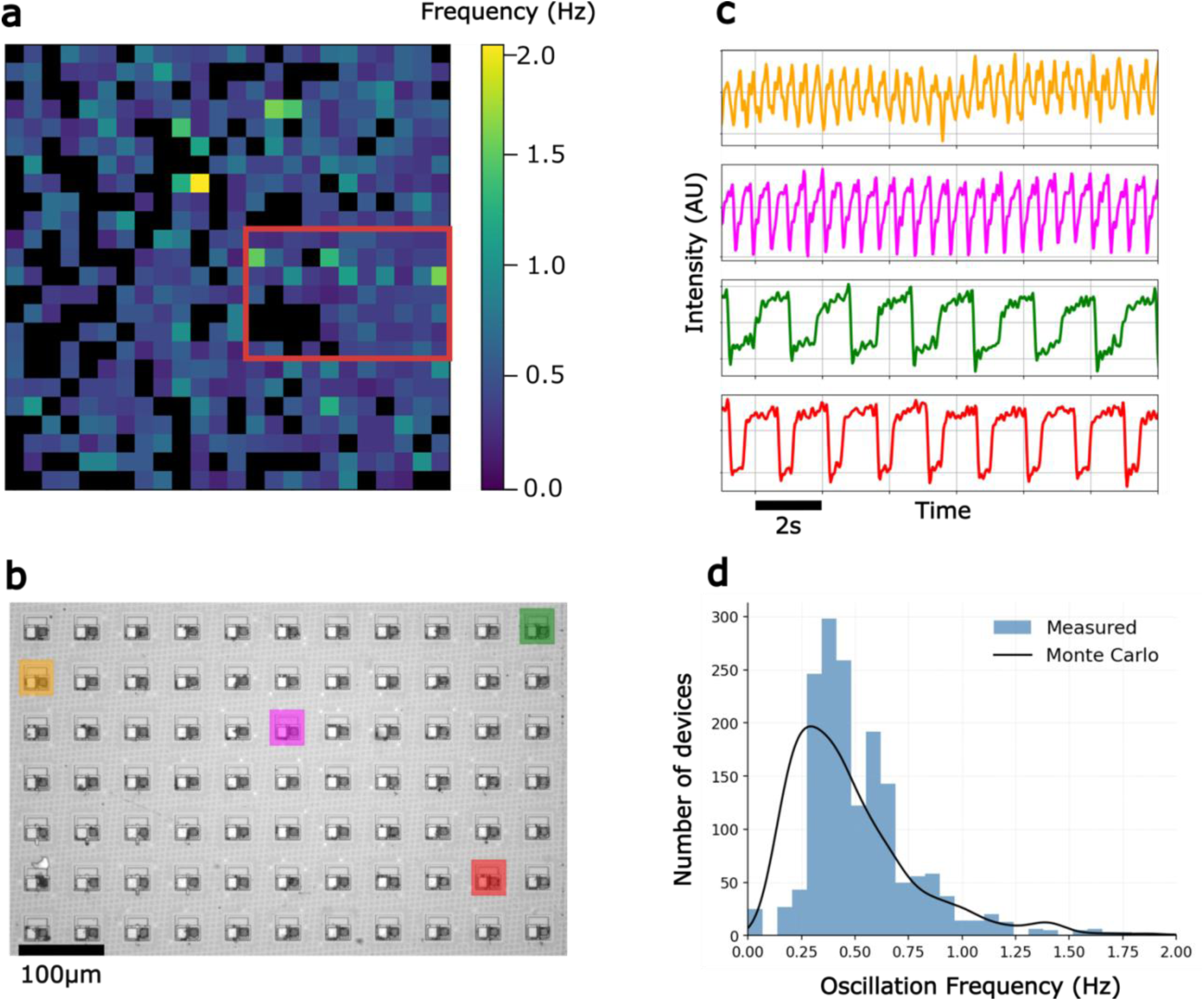
Benchtop optical testing of post-processed chips. **(a)** *f_oscillation_* map of 576 post-processed GLRO motes on a die before release from the CMOS substrate. Black squares indicate non-operational motes (due to yield issues with the modulator fabrication). **(b)** Brightfield image with selected annotated motes with temporal responses shown in **(c)**. **(d)** Histogram of measured *f_oscillation_* across 1657 operational motes tested across five CMOS dies. The solid line is the predicted variability from Monte Carlo spice simulations. All the data shown here is from the 30µm GLRO motes.

### Temperature sensing

The electrical behavior of the motes is sensitive to temperature due to the underlying temperature sensitivity of the silicon electronics. **Fig. 5a** and **Fig 5b** highlight the temperature sensitive components of the RO delay cell and the GLRO mote respectively. There are two components of the mote circuitry that are responsible for its temperature sensitivity: the photodiode and the oscillator circuitry. The photodiode modulates *f_oscillation_* by inducing a drop in the supply voltage with increasing temperature. The transistors that make up the mote circuitry turn on more easily with increasing temperature and this affects *f_oscillation_* as well. These two effects are in contention in both the RO and GLRO mote. They largely cancel out in the GLRO mote, but leave a residual negative temperature sensitivity in the RO mote. A more detailed discussion of the mote temperature sensitivity is provided in the **Supplemental Text**.

**Figure 5.**
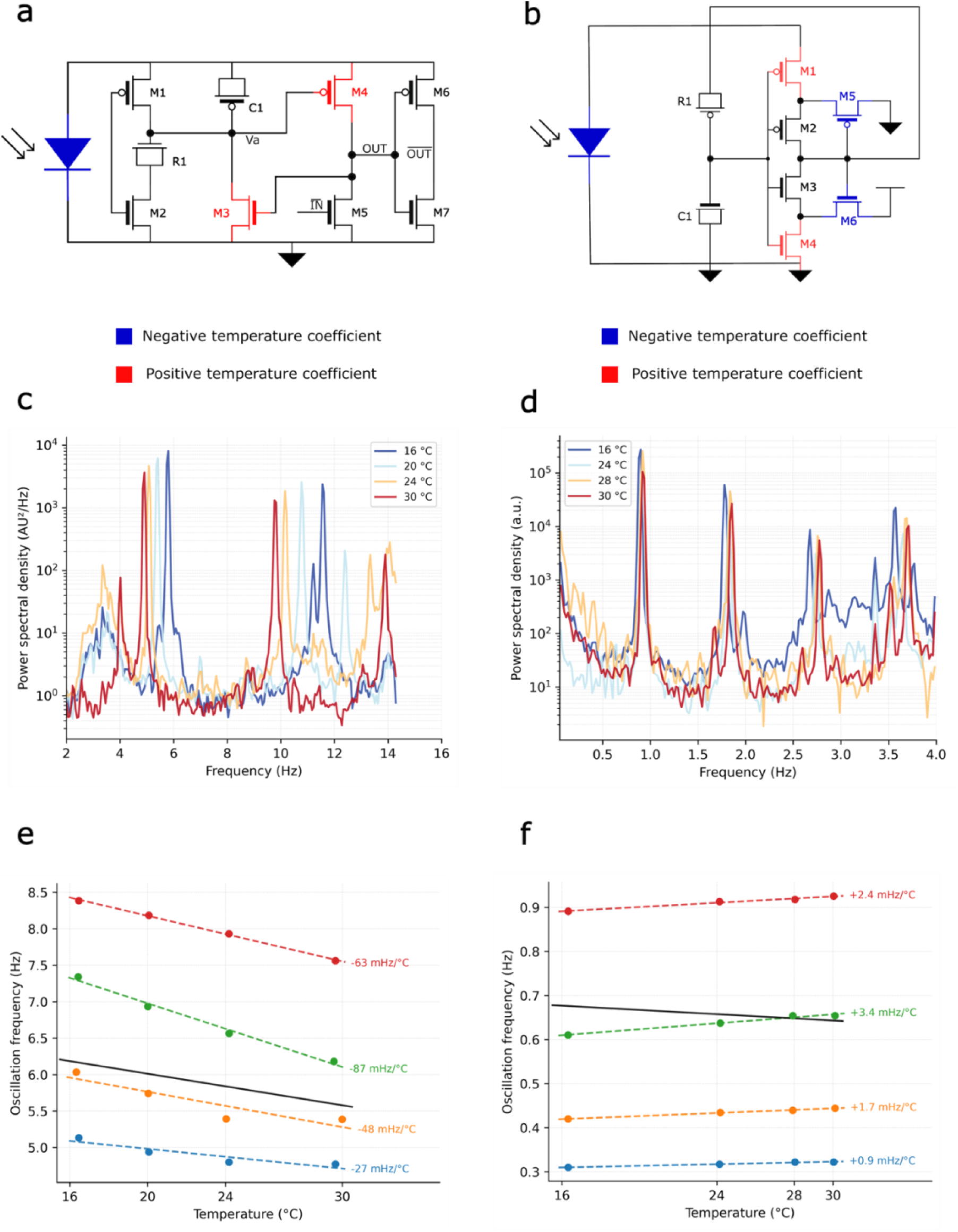
Temperature sensitivity characterization of motes. Temperature sensitive components of the circuit highlighted in **(a)** Ring oscillator (RO) delay cell and **(b)** Gate-leakage relaxation oscillator (GLRO). *f_oscillation_* as a function of temperature for RO and GLRO motes is shown in figures **(c)** and **(d)**. Each dashed curve is a different measured device The solid black curves are the predicted simulated temperature sensitivities of nominal motes. Power spectral density of a single RO **(e)** and GLRO **(f)** mote captured at different bath temperatures.

We evaluated the temperature sensitivity of the motes at scale before their release into solution. The motes were measured in a regulated temperature environment in a water-jacket-cooled microscope-stage-top incubator system. The illumination power matches that used in the experiments of **Fig. 4** (was 8 nW/μm^2^), and the same baseline removal and denoising strategy was employed. A power spectral density (PSD) was constructed from a 108-second-long capture time trace with the optical image being sampled at a rate of 35 frames per second (the maximum supported frame rate for a full ROI capture) and the fundamental frequency was identified. As shown in **Fig. 5c,d**, the fundamental frequency changes with temperature with a negative sensitivity for the RO and a weakly positive sensitivity for the GLRO. Multiple measured motes all displayed a similar behavior although the sensitivity varied across measured motes (**Fig. 5e,f**) but matched predictions from circuit simulation as shown with the solid black curves in **Fig. 5e,f**. The variability in temperature sensitivity can be attributed to the highly sensitive gate-leakage-based design, requiring per-mote calibration. The average temperature sensitivity of the RO mote is close to -70 mHz/°C with a 1-σ spread of 50% over 71 measured devices (**Fig. S6b**) while the GLRO mote is quite insensitive to temperature across measured devices (**Fig. S6a**). Since the frame rate is approximately 35 frames per second, the Nyquist frequency is 14.2 Hz and third harmonic content from the RO mote aliases back in-band at the camera acquisition rate. The higher frequency components in the PSD (**Fig. 5c,d**) are the harmonics of the fundamental in the Nyquist and aliased bands. Encoding the temperature reading in the frequency of the telemetry response (rather than its amplitude) makes the readout insensitive to confounding effects such as the temperature sensitivity of the amplitude of the QCSE modulator.

### Testing of the released motes

The motes were released in solution using the procedure outlined in **Fig. 3** and **fig. S7a** and imaged under widefield epifluorescence microscopy. Over 700 motes were released in solution at scale and transferred to a petri dish for imaging. Their extremely small size necessitated careful manipulation and handling under a microscope. We used pulled glass micropipettes with an inner diameter (ID) of approximately 20 µm under controlled suction to pick up and move motes, similar to approaches used to handle individual cells. Using this approach, operational motes could be identified, sorted, and individually transferred for imaging as shown in **fig. S7b**.

We observed that the overall yield drops to 5-10% post release because of losses in the release and handling processes. **Fig. 6a,b** shows multiple operational motes under fluorescent (upright) and white light (inverted) microscopy. Oscillatory fluorescent responses of selected motes (denoted in **Fig. 6a** as Motes 1-8) is shown in **Fig. 6c. Supplementary Movie S1** shows the oscillatory fluorescence of Motes 1, 4 and 7 from **Fig. 6a**.

**Figure 6.**
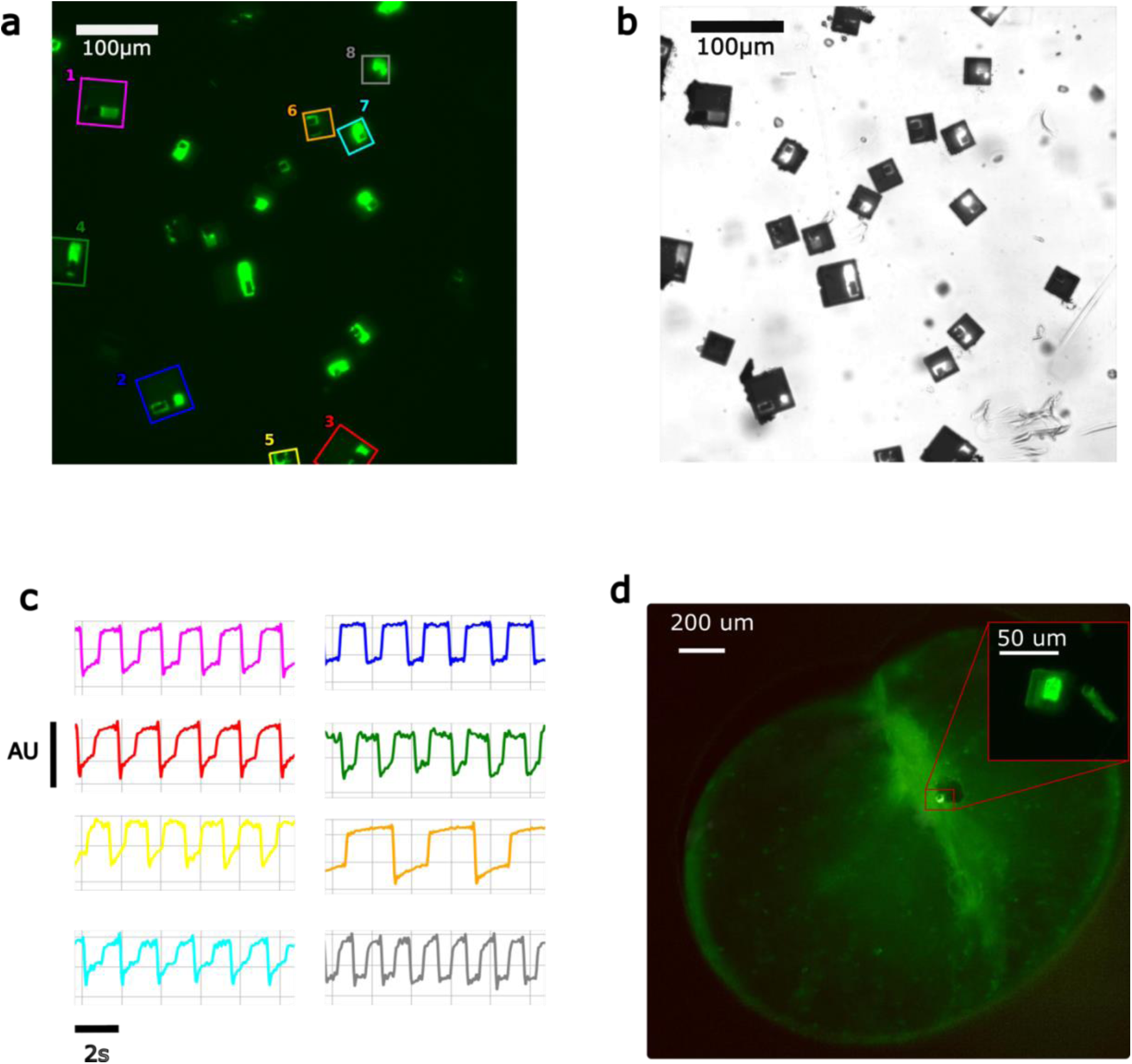
Optical measurements of released chips in vitro. **(a)** Epifluorescent image of released motes with eight operational motes highlighted. **(b)** Brightfield image of motes shown in (a) under diascopic illumination. Both 30μm and 50μm motes are visible. **(c)** Time trace of measured fluorescence intensity of the eight motes highlighted in (a). **(d)** Epifluorescent image of mote inside a cell of *Noctiluca scintillans*. The mote is a 30μm variant and the cell is ∼1.6 mm in diameter.

### Intracellular incorporation of the motes

*Noctiluca scintillans* (commonly known as Sea Sparkle for its bioluminescent glow) is a large green dinoflagellate that is a mixotroph that combines carbon-fixation from its autotrophic endosymbiont *Protoeuglena noctilucae* with phagocytotic ingestion of prey. These plankton can grow very large (200-1000 µm) and are mainly found in the Sea of Oman, the northern Arabian Sea and along the coasts of Southeast Asian countries. Since the turn of the millennium, blooms of *N. scintillans* have largely replaced diatom blooms in part because of their dual trophic modes of meeting their metabolic and growth needs and the superior autotrophic ability of their endosymbionts *P. noctilucae* in hypoxic waters (*28*). The rise in *N. scintillans* is already having an adverse impact on the food web in the Arabian sea because *N. scintillans* are not readily consumed by mesozooplankton that function as a crucial intermediate link in the food web between phytoplankton and larger aquatic predators. Because of their large size, the presence of endosymbionts within their central cytoplasm, their distinctive role in the ecology of the oceans and their phagocytotic capability, they are an ideal model organism for studying intracellular processes through incorporation of these motes.

We developed a method to implant these motes intracellularly into *N. scintillans*. The procedure (depicted in **Fig. S8e**) involves incubating starved populations of cells, deprived of both light and food, in f/2 growth media, while motes are transferred to the saline buffer. Starving the cells stimulates feeding behavior and enhancing mote uptake. To increase encounter probability, cells and motes were carefully drawn into the narrow stems of sterile transfer pipettes, prompting close physical interactions. During phagocytosis, *N. scintillans*, ingests the motes as if they were prey. The motes are initially enclosed in a food vacuole, whose outer membrane gradually ruptures releasing the chips in the cytosol. Despite enhancing the encounter probability, ingested mote yields were very low. An average of 10 motes and 5 cells were suspended in a volume of 2ml of f/2 media within each transfer pipette. 10 such pipettes were setup for incubation and only 2 cells with ingested motes were identified for an overall uptake yield of 2%. Further work needs to be done to improve the process yield of the mote fabrication and release as well as the yield of the uptake process. **Fig. 6d** shows a representative image of a cell containing a CMOS mote in its cytosol and **fig. S8c,d** shows representative images of cells containing passive dummy silicon chips of comparable size, incorporated through the same incubation process in initial testing of the incubation protocol. More details about the ingestion procedure can be found in the **Methods** and **fig. S8**.

## Discussion

Here we demonstrate a fully autonomous, optically powered CMOS mote with a volume below 10 pL with a sub–10 pW power envelope, which to our knowledge represents the smallest active intracellular-scale electronic system reported to date. By combining aggressive CMOS scaling, monolithic optical power harvesting, and fluorescence-based backscatter telemetry, this work extends active bioelectronics into the intracellular domain, previously accessible only via passive tags or molecular probes. More importantly the ability to fabricate, release, and optically interrogate thousands of such motes in parallel establishes a practical path toward scalable, cell-scale electronic interfaces.

A central contribution of this work is the use of a fluorescence-based backscatter modulator to achieve wireless telemetry under extreme power and volume constraints. Unlike light-emitting sources, which require substantial static and dynamic power, the capacitive QCSE modulator consumes negligible static power and leverages the energy of an external optical field for signal generation. This architectural choice reallocates the limited harvested energy budget toward on-chip computation and sensing, while enabling substantial miniaturization. Importantly, the fabrication flow supports the integration of multiple modulators on a single mote with a shared reference, enabling redundancy, increased data throughput, or spectrally multiplexed readout.

The use of quantum-dot–based fluorescence modulation further introduces flexibility in optical system design. Because emission wavelength is tunable through nanocrystal composition and size, motes with distinct emission spectra could be deployed simultaneously to support functional multiplexing or to coexist with conventional fluorescent reporters. In addition, information can be encoded in multiple domains, including frequency, duty cycle, or amplitude and allowing the telemetry strategy to be adapted to different sensing modalities or imaging constraints. Encoding information in the frequency domain, as demonstrated here for temperature sensing, also renders the readout robust to variations in excitation intensity, photobleaching, and temperature-dependent fluorescence efficiency of the modulator itself.

The optical nature of both power delivery and telemetry introduces constraints that are intrinsic to this operating regime. The orientation of the mote relative to the excitation field affects both harvested power and emission efficiency, necessitating consideration of optical access and imaging geometry. While these effects can be partially mitigated using upright and inverted microscopy or by engineering more isotropic photodiode and modulator geometries, they highlight a fundamental tradeoff between miniaturization and orientation sensitivity. Additionally, interference effects from the CMOS backend stack impose wavelength-dependent filtering, which must be accounted for when selecting excitation and emission bands. These limitations are not unique to this system but are likely to be common to future intracellular optoelectronic devices fabricated in standard CMOS processes.

Once released into solution, motes can be manipulated and positioned using established micropipette-based techniques, enabling targeted placement for localized sensing experiments. While the present work focuses on temperature sensing as a proof-of-concept, the use of standard CMOS technology greatly simplifies the integration of additional sensing modalities, including pH, ion concentration, or electrophysiological measurements, as well as more complex on-chip signal processing.

Intracellular delivery of motes into *N. scintillans* demonstrates the feasibility of embedding active electronic systems directly within the cytosol of a living cell with minimal volumetric displacement. The delivery yield (2%), multiplied against with the yield of the fabrication and release process (5-10%), make delivering functional motes into cells incredible difficult at the fabrication scale of hundreds of motes. Future work will need to address scale, delivery efficiency, and fabrication yield to drive these techniques to applications.

This work points toward a new class of bioelectronic tools that complement existing molecular, genetic, and optical approaches. Intracellular CMOS motes could enable continuous, closed-loop interrogation of cellular physiology, bridging fast electrical dynamics with long-term biochemical state changes. By operating at power densities comparable to cellular metabolic rates and at volumes compatible with cytosolic integration, such devices open the possibility of persistent electronic sensing and actuation at the single-cell level. As CMOS scaling and low-power circuit techniques continue to advance, intracellular bioelectronics may evolve from single-function motes to programmable, networked systems capable of interacting with living cells over extended timescales.

## MATERIALS AND METHODS

### Modulator microfabrication

A 15mm x 15mm silicon substrate is prepared by creating a 3.5 mm x 3.5mm x 300μm recessed pocket on a planar silicon die using a Bosch process. The CMOS chip is bonded face-up into the recessed pocket using Norland Optical Adhesive (NOA 86H) and cured under heat and pressure (125 C, 5N, 10min) using a Fineplacer Lambda die bonder. This process offers a planar surface near the edges to reduce edge-beading when working with spin-coated materials and the substrate also plays the role of a handle die. This is the starting point for the microfabrication process.

First, the CMOS passivation dielectric is etched in CF_4_ plasma using an ICP-RIE plasma process (CF_4_, 10 mTorr, 1000 W ICP, 20 W RIE) to remove the top 1800 nm of the Si_3_N_4_/SiO_2_ dielectric passivation until the top layer of copper pads are exposed. The surface is inspected and optionally a second process of slow wet etching in BOE (buffered oxide etchant) 10:1 is employed to expose the pads. The sample is then transferred to a vacuum chamber for deposition of a metallic layer to interface with the modulator. The surface is first pre-cleaned by sputtering in an Argon environment and 50 nm/100 nm of Cr/Al_99_Si_1_ is deposited. A photoresist etch mask is used to define the pads on a 0.5μm thick layer of S1805 (Kayaku Advanced Materials) exposed using a Direct-write laser lithography system (Heidelberg DWL 66+) developed in a Tetramethylammonium hydroxide (TMAH) based developer (AZ 300 MIF). Finally, the Cr/AlSi layer is patterned by etching through the openings of the photomask in a chlorine plasma (3:4 Cl_2_:BCl_3_, 20 mTorr, 1200W ICP, 100W RIE) and the residual resist is removed. This concludes the definition of the metal pads (**Fig. 2d**).

The next step involves the deposition and patterning of the modulator. The modulator stack is defined using three consecutive deposition steps of HfO_2_, quantum dots and HfO_2_. The bottom and top hafnia layers, each 10 nm thick are deposited using thermal ALD processes at 200 °C with a TMDAH (Tetrakis(dimethylamido)hafnium) precursor. The sandwiched quantum dot layer is deposited by spin coating a controlled volume (45 μL) of a 2.2 nmol/ml aqueous solution of colloidal green quantum dots (HECZW520, NN Labs) at 300 rpm for 15 minutes. The entire HfO_2_/QD/HfO_2_ stack is around 150 nm thick. After the top hafnia layer is deposited, the photoactive modulator regions are defined using a 1.8 μm photoresist mask (S1818 / DWL 66+ / AZ300 MIF). Finally, the modulator is patterned by etching the hafnia and the quantum dot in a chlorine plasma (3:4 Cl_2_:BCl_3_, 20 mTorr, 1200W ICP, 100W RIE) and excess resist is stripped out. This stage of the process is highlighted in **Fig. 2e**.

Finally, the top ITO layer is deposited via DC sputtering from an ITO target in an Ar atmosphere (5 mTorr, 180W) yielding a 100 nm thick film. This film is then etched under a photoresist mask (S1818 / DWL 66+ / AZ300 MIF) in a CH_4_/H_2_/Ar plasma (5:2:2, CH_4_:H_2_:Ar, 10 mTorr, 1000W ICP, 200W RIE). The excess resist is stripped (**Fig. 2f**), and the entire die is passivated with 1-μm-thick PECVD SiO_2_ deposited at 150 °C. At this point the die is tested for modulator function before the release process.

### Mote release

A trench etch mask is defined and patterned with a combined metal/photoresist mask (Cr 200nm / AZ 10XT, 8um). Trenches are etched into the backend oxide stack using a combination of wet etch (BOE 10:1) and plasma etch (CF_4_ ICP-RIE). The trench is then etched down into the silicon using a Bosch etch process until the trenches are around 12 μm deep into the substrate.

The residual resist is stripped out and a passivation stack of 200-nm-thick SiO_2_ (PECVD). The die is then bonded facedown onto a glass handle wafer with CrystalBond-509 allowing it to creep into the crevices and hold the unreleased motes in place (**Fig. 3d**). The die is then flipped over, and the silicon pocket substrate and the CMOS die are removed from the backside (ICP, SF_6_ plasma) until the motes are singulated under visual inspection. At this point in the process, the 10um thick Si motes are embedded in CrystalBond but still held together by the SiO_2_ passivation which is resilient to the selective chemical etch (**Fig. 3e**). The 200 nm-thick SiO_2_ passivation is dry-etched using ICP-RIE (CHF_3_, 30 mTorr, 200 W ICP, 30 W RIE). Finally, the motes are released from the CrystalBond matrix in acetone, centrifuged and resuspended in a proprietary NMP based solvent (Remover PG), and are stored long-term in the dark at a temperature of 4 ℃.

### Mote handling

The motes are handled and transported in solution in a 35-mm glass petri dish under diascopic white light microscopy. Manipulation of the motes is accomplished using manually pulled (P-87 micropipette puller, Sutter Instruments) glass pipettes that are beveled to have an inner diameter between 15 μm and 25 μm. The glass pipettes are attached to a holder mounted on a three-axis micromanipulator and are connected to a regulated suction controller. Suction-based control is then used to pick and place the motes in solution and to orient them appropriately for imaging.

### Mote imaging and detection

The motes are imaged under blue light in a BX51WI microscope and the fluorescence emission is captured through a green bandpass color filter. Fluorescence images are captured using a Hamamatsu scientific camera with an exposure time of 50ms. To detect working released motes, the camera image is processed by detecting and removing the image background and computing the FFT and the SNR of the image content under a rolling mask over the entire image. The SNR map is then used to segment the image to aggregate adjacent candidate pixels and classify them as belonging to the same working mote.

### Ingestion of motes by Noctiluca

*N. scintillans* cultures used in this study were originally isolated from open waters of the Arabian Sea. The cultures were maintained in f/20 medium without silicate in polycarbonate bottles at 26 °C under an irradiance of 200 µE m⁻² s⁻¹ with an 11.5-h-light /12.5-h-dark photoperiod. As a mixotrophic dinoflagellate, *N. scintillans* can be sustained either through provision of external prey or maintained without supplemental prey under controlled laboratory conditions.

To evaluate the capacity of *N. scintillans* to ingest engineered microdevices, we first validated ingestion behavior using fluorescent tracer particles. Individual *N. scintillans* cells were gently isolated and transferred into sterile, filter-sterilized f/2 medium distributed in 24-well plates. Cells were pre-incubated for 48 h in darkness to stimulate feeding activity. Following this starvation period, cells were exposed to amine-modified polystyrene fluorescent latex beads (Sigma-Aldrich) spanning diameters of 10–250 µm and returned to controlled incubation conditions. Fluorescence microscopy performed after overnight incubation confirmed active ingestion across the tested size range, including beads up to 250 µm in diameter (**fig. S8a**), demonstrating the capacity of *Noctiluca* to internalize large particulate material.

We subsequently tested ingestion of biogenic silica particles (∼150-µm-diameter) using the same well-plate configuration. No uptake was observed after 7 days. However, detailed inspection revealed that, unlike the buoyant latex beads, silica particles rapidly settled to the bottom of the wells, limiting encounter rates with the vertically motile *Noctiluca* cells (**fig. S8b**)

To overcome this physical limitation, we employed sterile transfer pipettes to intermittently resuspend both cells and particles within the wells. Gentle bulb compression facilitated redistribution of settled particles into the water column, increasing particle–cell contact probability. Under these modified conditions, experiments with 150-µm-sized inert “dummy” silicon chips yielded successful ingestion events (**fig. S8c, d**).

Encouraged by these results, we proceeded to test functional microelectronic chips (30–50 µm diameter). Experimental protocols mirrored those used for bead and dummy-chip trials. Individual cells were monitored microscopically to confirm internalization.

## Funding

This work was supported by the:

Gordon and Betty Moore Foundation (KLS, JG)

Chen-Zuckerberg Biohub Network (KLS)

## Author contributions

Conceptualization: GR, KLS

Methodology: GR, FC

Investigation: GR, FC, WS, VA-P,

Visualization: GR

Supervision: KLS, JG

Writing—original draft: GR, KLS

Writing—review & editing: GR, FC, KLS

## Competing interests

GR and KLS are listed as inventors on a patent filed by Columbia University (patent no. US 12,247,919 B2, published 11 Mar 2025). The other authors declare that they have no competing interests.

## Data and materials availability

All data needed to evaluate the results in this paper are available in the main text or the supplementary materials.

## Supplementary Materials

### CMOS circuit design and operation

Two mote variants with different oscillators were designed as shown in **fig. S2**. Both oscillators employ gate leakage to achieve large time constants in an area-efficient fashion. Both variants were designed into motes of size 30 µm and the first variant was also designed into motes of size 50 µm.

The first variant, the gate-leakage-based relaxation oscillator (GLRO), shown in **fig S2a** employs a gate leakage element (R1) in the feedback loop of a Schmitt-trigger-based relaxation oscillator. The time constant is set by R1 and C1. Transistors M5 and M6 are configured to positively feed the output of S1 back into its input configuring it as a Schmitt trigger. The hysteretic thresholds can be tuned by tuning the relative sizes of the feedback devices M5, M6 and the pull up/pull down devices M1, M4.

The second mote variant, the gate-leakage-based three-stage ring oscillator (RO), implements a differential three-stage ring oscillator with a current starved delay stage shown in **fig. S2b.** The delay stages implement a gate-leakage based current starved element (R1) which sets the time constant along with capacitor C1. Transistors M3 and M4 constitute a regenerative latch which assists in propagating the input to the output after a fixed RC time constant.

### Spectral behavior of CMOS backend stack

As shown in **fig. S4**, the CMOS backend stack is made up of alternating thin films of materials with different refractive indices. The thickness of these films is often comparable to the wavelength of visible light in the material. As a result, the stackup ends up being spectrally selective and only light of certain wavelengths is allowed to reach the photodiode in the silicon frontend and the rest of it is reflected back. **Fig. S4a** shows the measured and simulated transmittance of the CMOS stack. The measurement was performed using thin film reflectometry – A beam of narrowband light is incident from a source on the sample and the spectrum of the light is swept while the reflected light is simultaneously captured. The light that is not captured as reflected is transmitted. This effect can be simulated if the wavelength-dependent refractive indices and thicknesses of the constituent stack layers are known with the transfer matrix method (TMM). The simulated plot in **Fig. S4a** is from a TMM simulation in which the known thicknesses and material properties of the stack were used to solve for the stack reflectance with a constrained local optimization performed on the thicknesses to fit to the measured curves. The constrained local optimization attempts to account for process variability in the stack layer thicknesses compared to the nominal values. The stackup derived from this fit is shown in **Fig S4b**.

### Temperature sensitivity of the CMOS motes

The motes derive their temperature sensitive properties both from the temperature-sensitive response of the photodiode as well as from the transistors that make up the oscillator. The photodiode’s sensitivity has a well understood negative coefficient with temperature. This reduces the supply voltage of the mote with increasing temperature.

The RO delay cell shown in **Fig 5a** has a strong positive sensitivity to temperature resulting from the action of the transistors in the regenerative latch (M3, M4). Both of these devices turn on faster at higher temperature and the delay of the delay cell is strongly tied to how fast the latch engages. At the same time, the sensitivity of its oscillatory response to supply voltage is also strong; as a result, the negative sensitivity resulting fro the influence of the photodiode on the supply voltage dominates. Both simulated and measured responses suggest a strong negative temperature coefficient of close to -60mHz/℃ (**Fig. 5e**).

The GLRO (**Fig. 5b**) also sees the influence of the photodiode and the influence of the core transistors in the circuit, but the hysteretic behavior of the Schmitt trigger (devices M1-M6) is fundamentally enabled by contention between M1, M5 and M4, M6 in the device. Making M1, M4 stronger makes the switching faster while making M5, M6 stronger makes the switching slower. This internal contention between similar devices in the circuit means that neither dominate over the other significantly with changes in temperature or supply voltage because they both see the same changes in drive strength. This leads to a lower sensitivity to temperature and supply voltage for the GLRO oscillator as shown in **Fig. 5f**. The GLRO is nearly 30 times less sensitive than the RO. Because of the flat response and the sensitivity of this internal contention to variability, the polarity of the simulated sensitivities do not always agree with measurements.

**Fig. S1.**
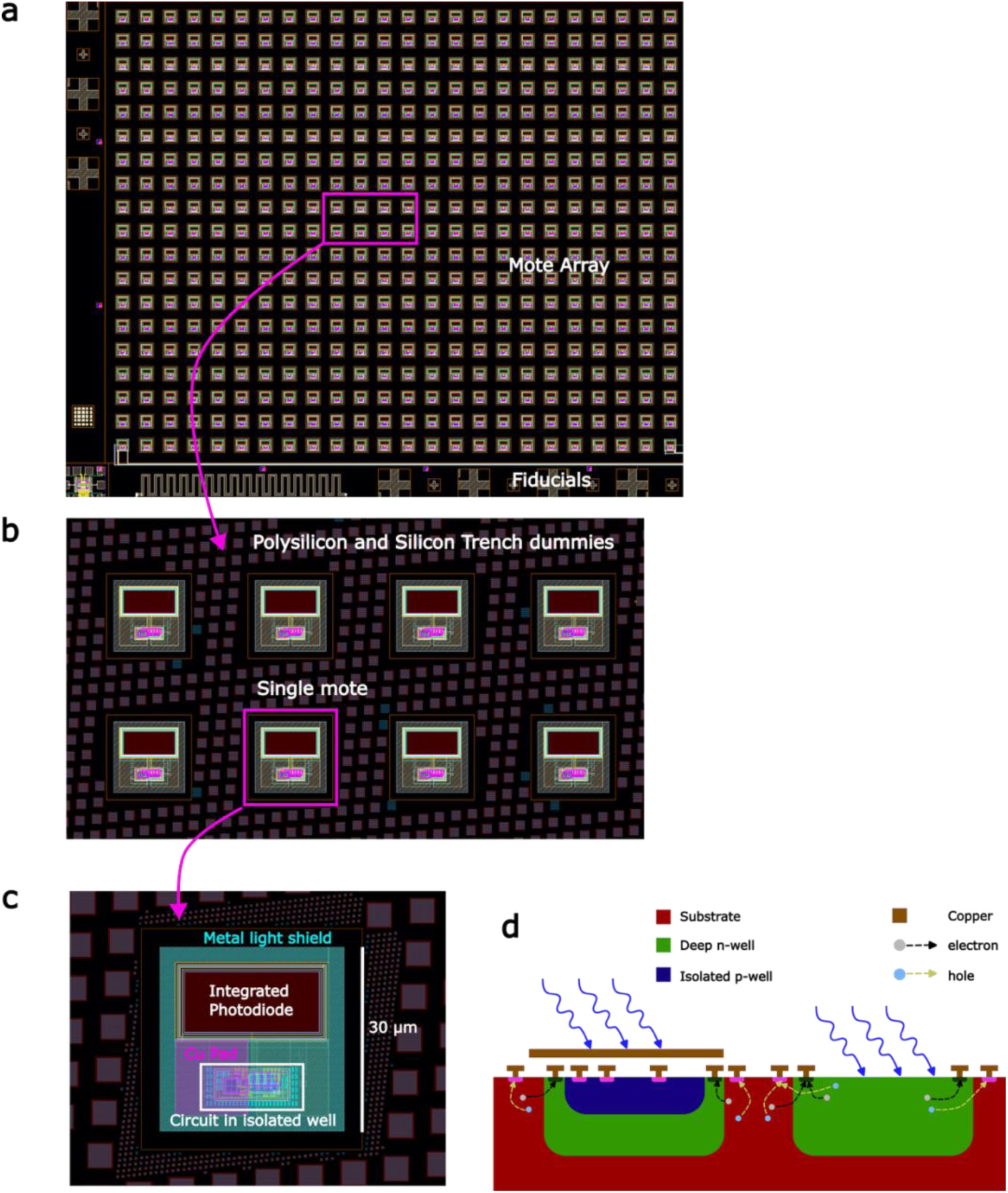
Layout of the CMOS motes. **(a)** Layout of the full die with motes separated by kerf regions for singulation. **(b)** Detailed view of several motes separated by kerf regions. **(c)** Single 30-μm mote layout. **(d)** Cross section of the mote. An integrated photodiode (right) is used for optical energy harvesting. The CMOS silicon circuitry is laid out in an isolated p-well (left) and covered by metallization to protect it from stray photocarriers from light and diffusing from the substrate.

**Fig S2.**
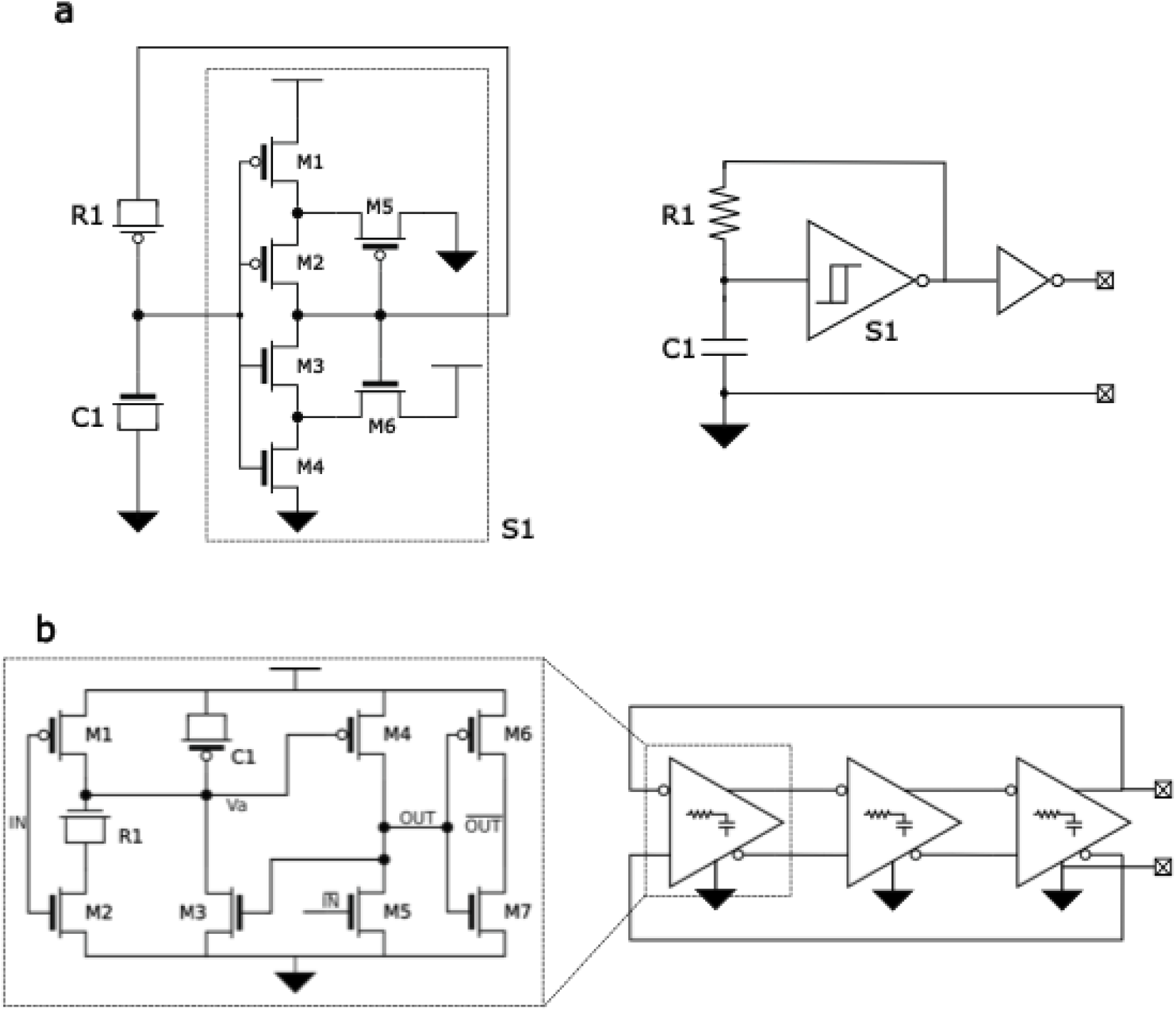
Two different variants of the temperature-dependent oscillator circuits employed. **(a)** Gate-leakage based relaxation oscillator (GLRO). **(b)** Gate-leakage based three stage current starved Ring oscillator (RO)

**Fig. S3.**
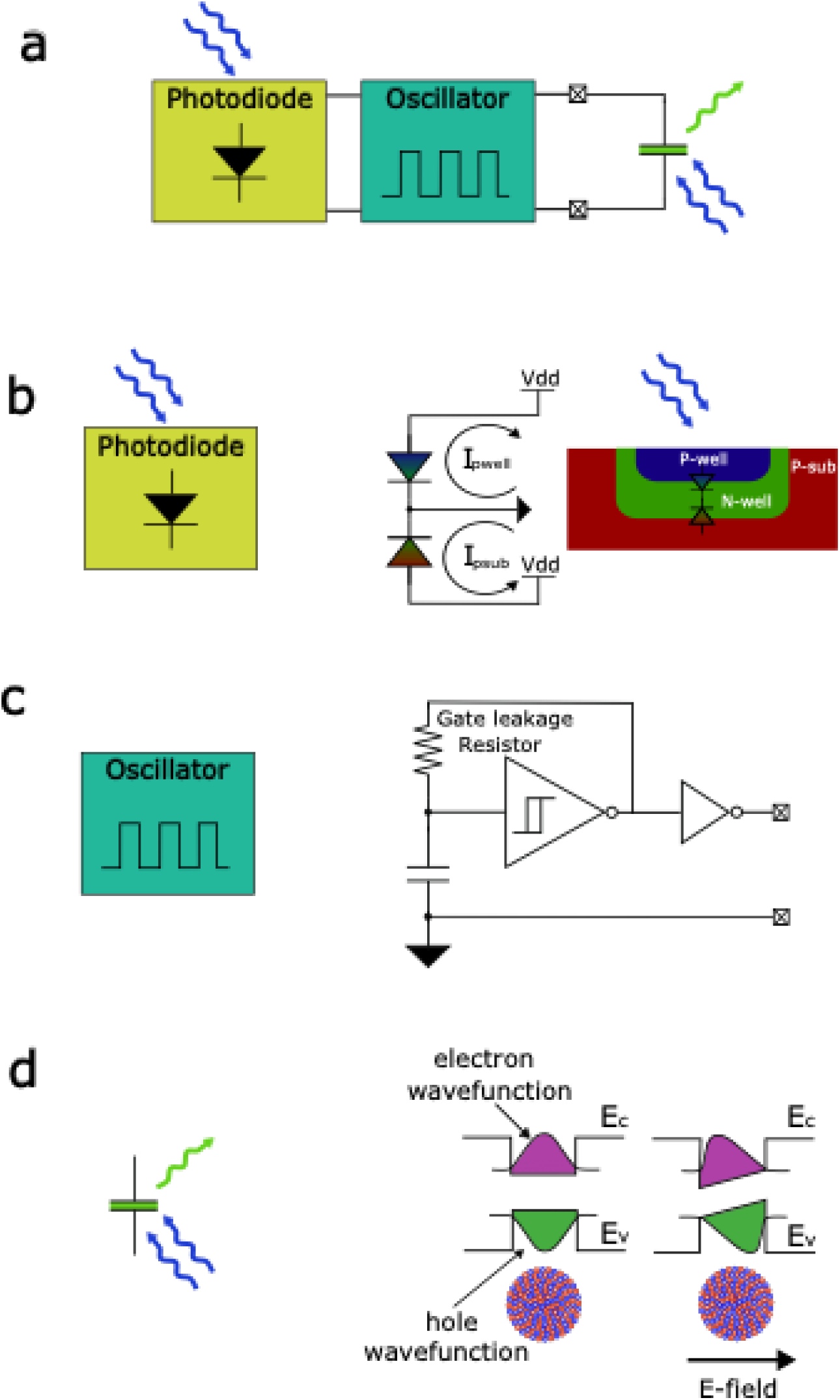
Constituent parts of the mote. **(a)** Block diagram with photodiode, oscillator and QCSE modulator. **(b)** Photodiode structure with multiple junctions participating in photon capture. **(c)** Relaxation oscillator implemented using gate leakage element. **(d)** Operation of the quantum confined stark effect modulator demonstrating electron-hole wavefunction overlap integral modulation due to band bending under the influence of an external electric field,

**Fig. S4.**
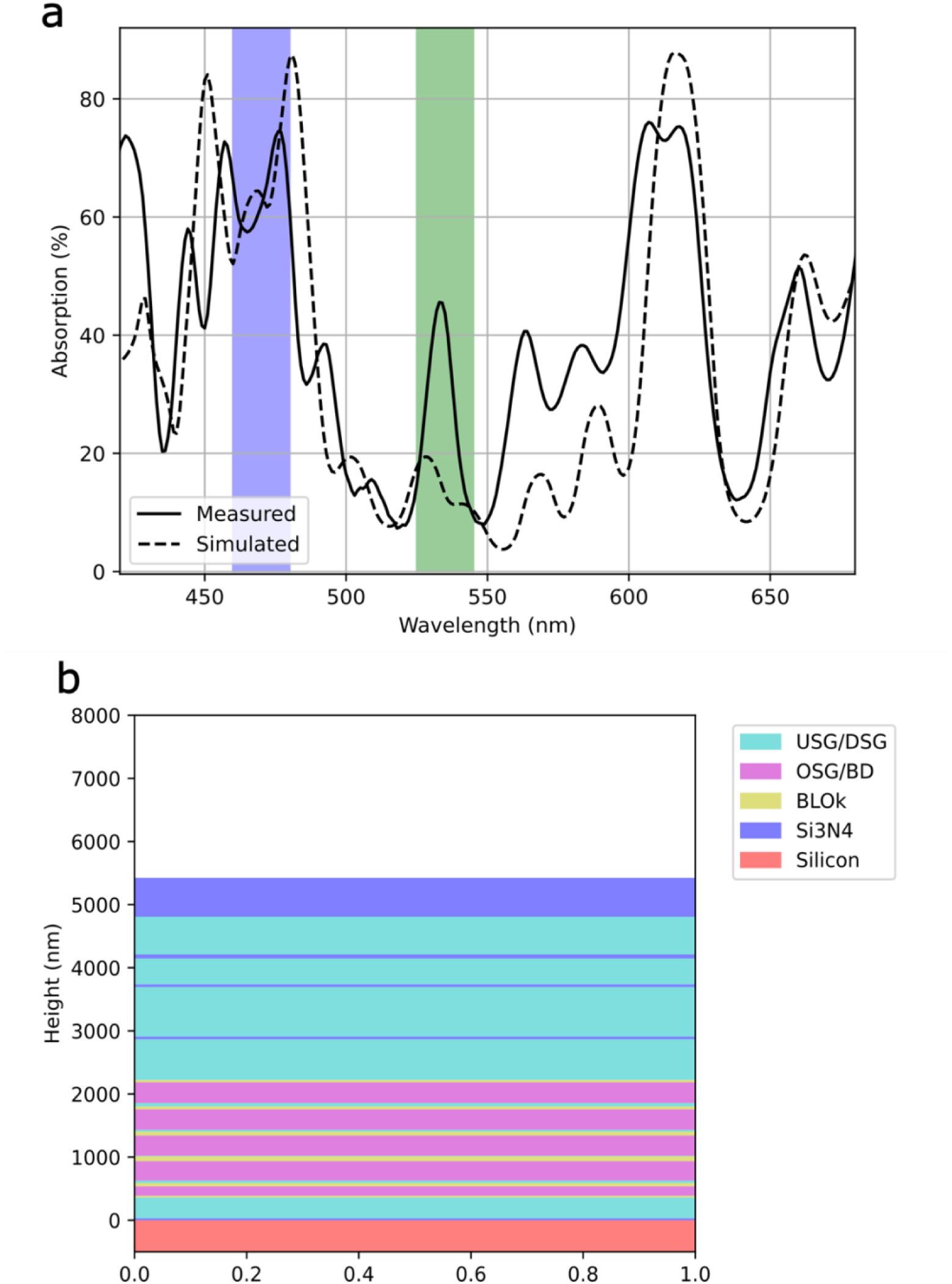
Simulation and measurement of the thin film spectral reflectance properties of the CMOS dielectric backend stack. **(a)** Excitation spectrum from the microscope and the emission from the quantum dots are annotated in blue and green. The thin film nature of the CMOS backend stack operates as an interference filter and confers a strongly wavelength-dependent quantum efficiency to the CMOS photodiodes. **(b)** Illustration of the backend dielectric stack of the 6 metal 65nm bulk CMOS process highlighting the oxide, nitride and low-k dielectric layers.

**Fig. S5.**
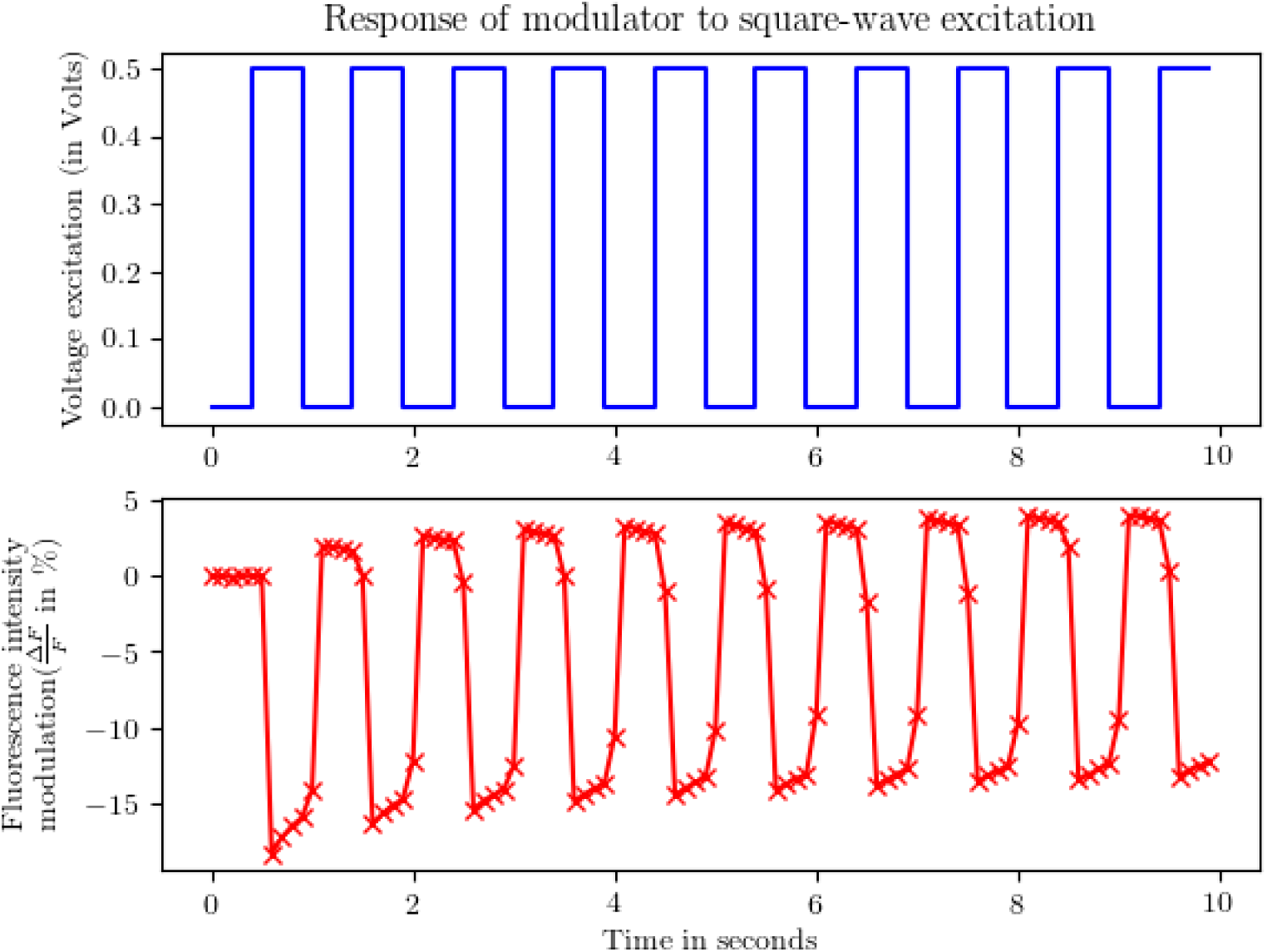
Characterization of QCSE modulator function. Voltage waveform applied across standalone modulator terminals (top) and sampled measurement of change in fluorescence intensity with applied voltage (bottom). Measurement was performed on a standalone QCSE modulator actuated by an external voltage source.

**Fig. S6.**
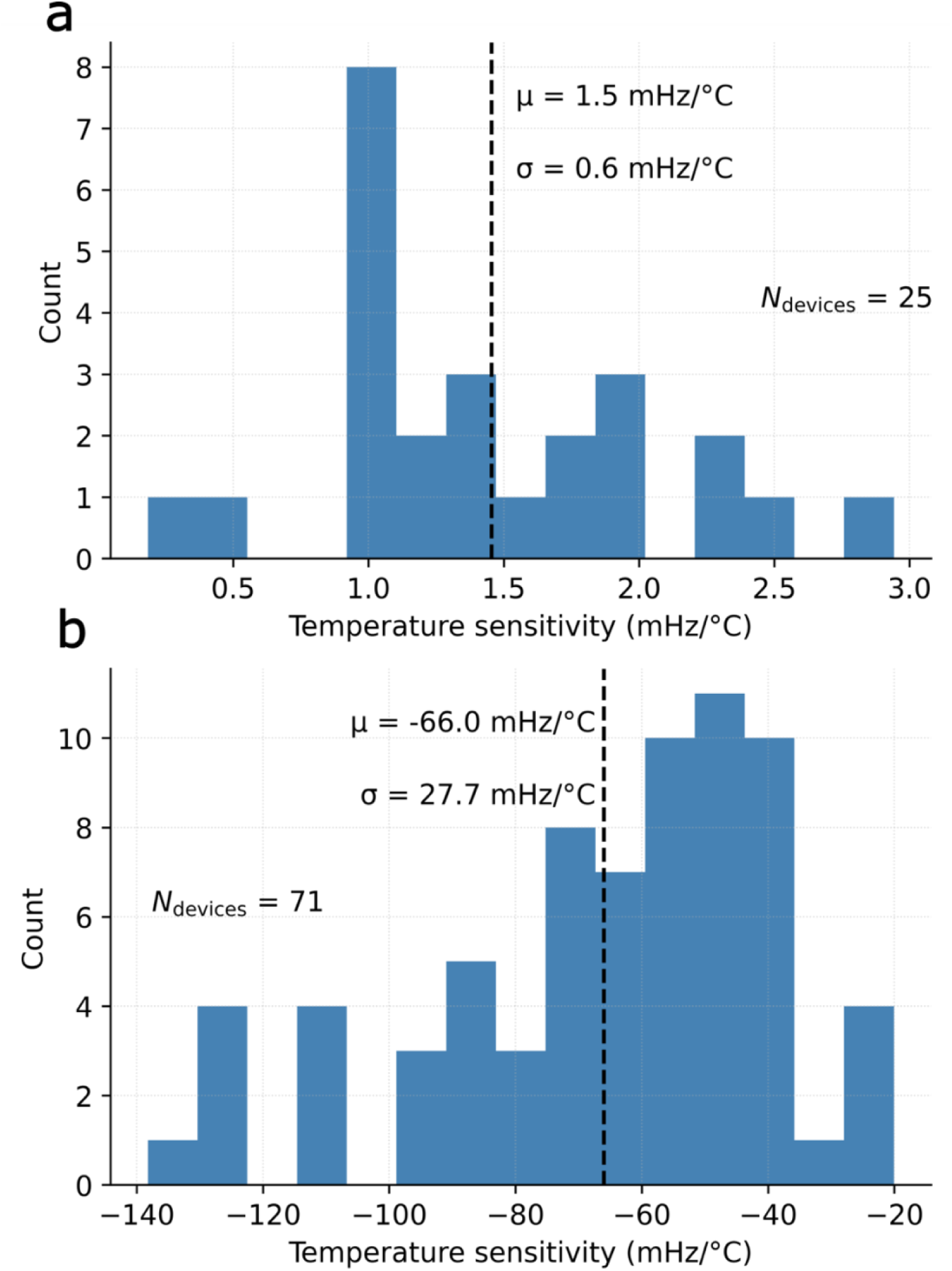
Measured temperature sensitivity of the motes. (a) Temperature sensitivity across 25 measured GLRO motes. The GLRO exhibits weak positive temperature sensitivity. (b) Temperature sensitivity across 71 measured RO motes. The RO motes exhibit stronger negative temperature sensitivity.

**Fig. S7.**
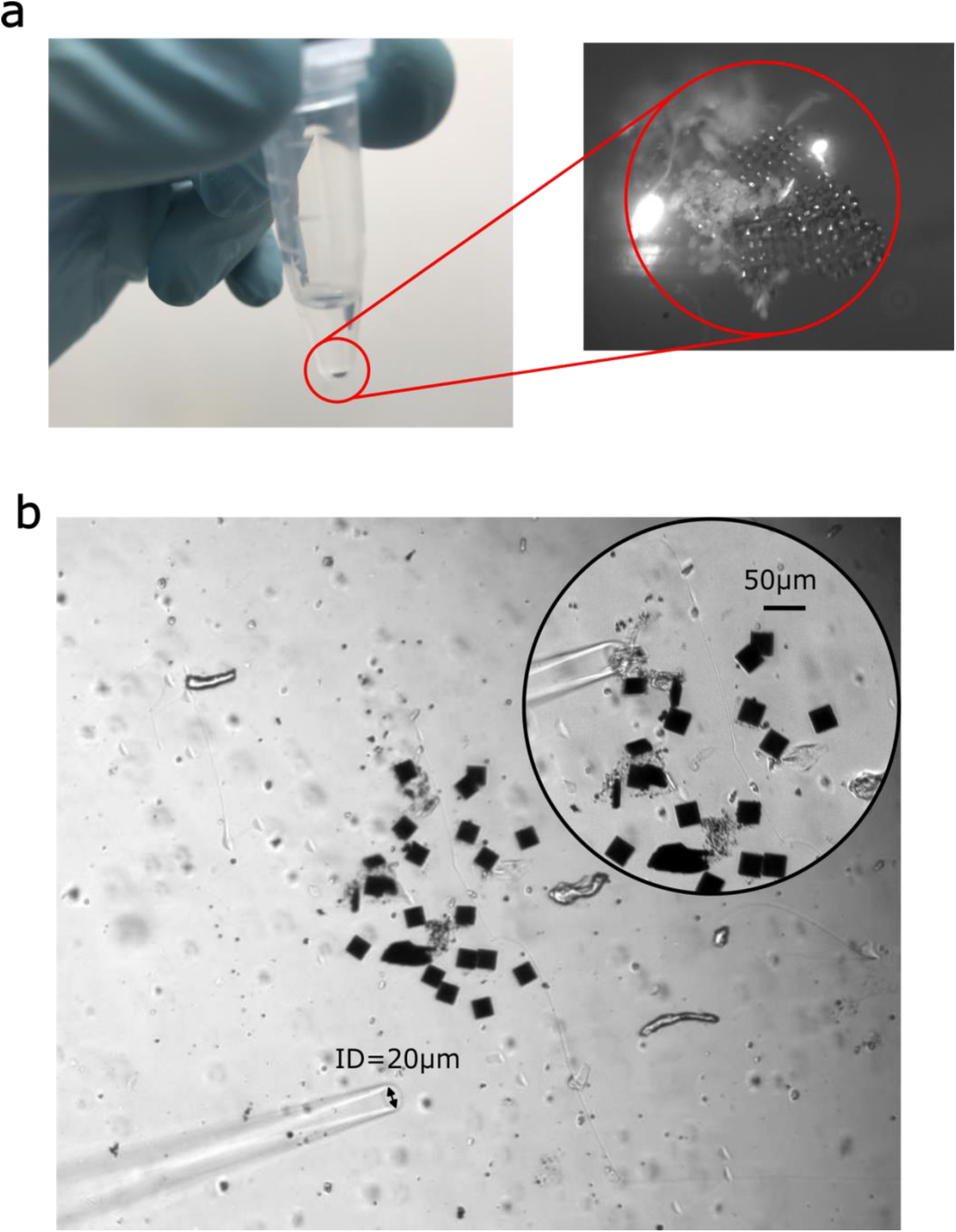
Release and manipulation of motes. **(a)** Pellet of released motes suspended in the bottom of an Eppendorf tube after release. Epifluorescent microscope image of pellet in the inset. The motes are still held together by some CrystalBond prior to release by ultrasonication. **(b)** Micropipette used to transfer the motes in solution. Image shows motes collected together using the suction-based manipulation using the pipette. Inset shows motes and pipette in close proximity.

**Fig. S8.**
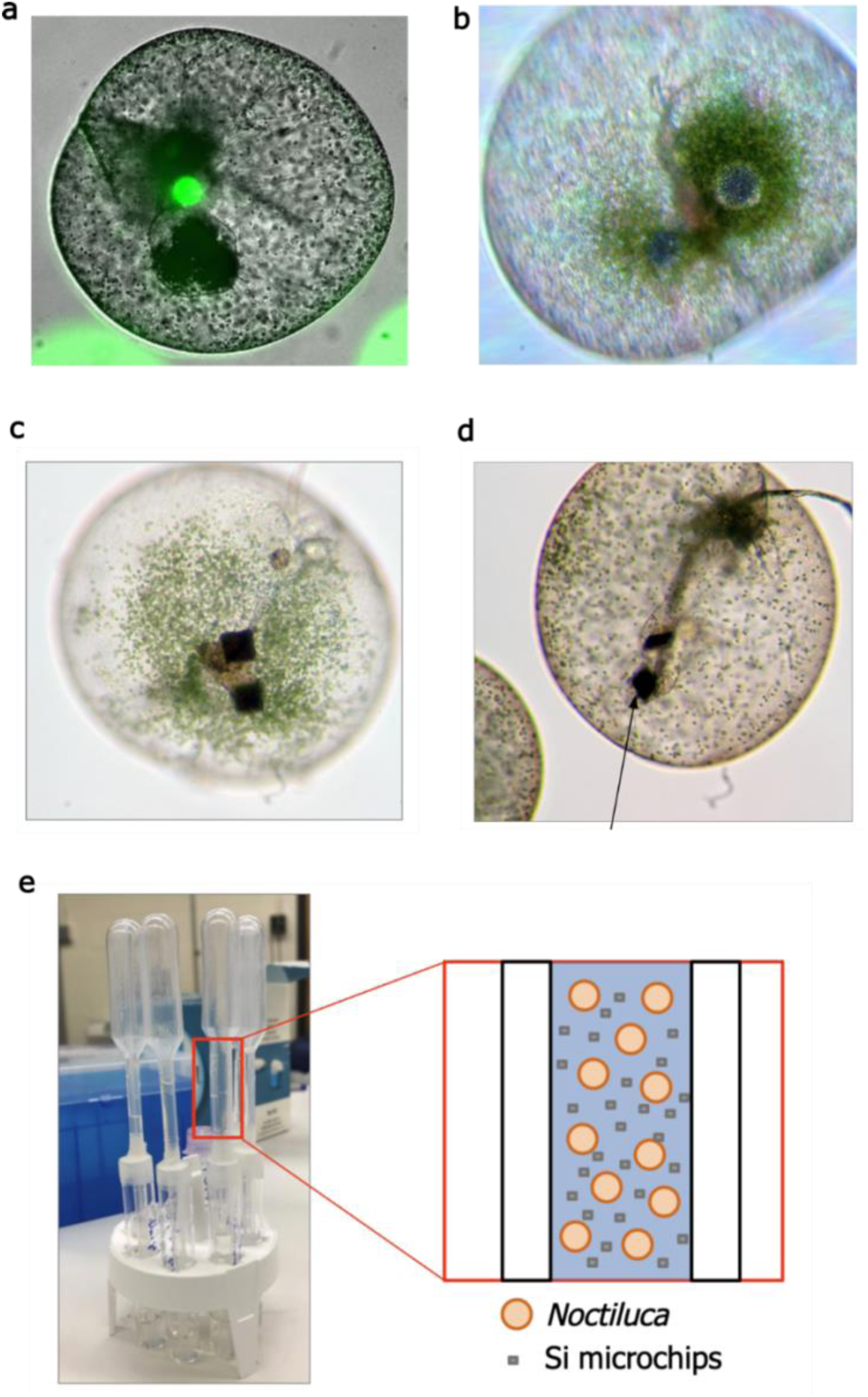
Incorporation of chips into *N. scintillans*. **(a)** *N. scintillans* with ingested fluorescent beads. **(b)** *N. scintillans* with ingested silica particles. **(c), (d)**. 150 μm × 150 μm × 150 μm dummy silicon chips ingested by *N. scintillans* using the setup outlined in (e). (e) Setup to suspend noctiluca and chips in the neck of a pipette.

**Movie S1.**
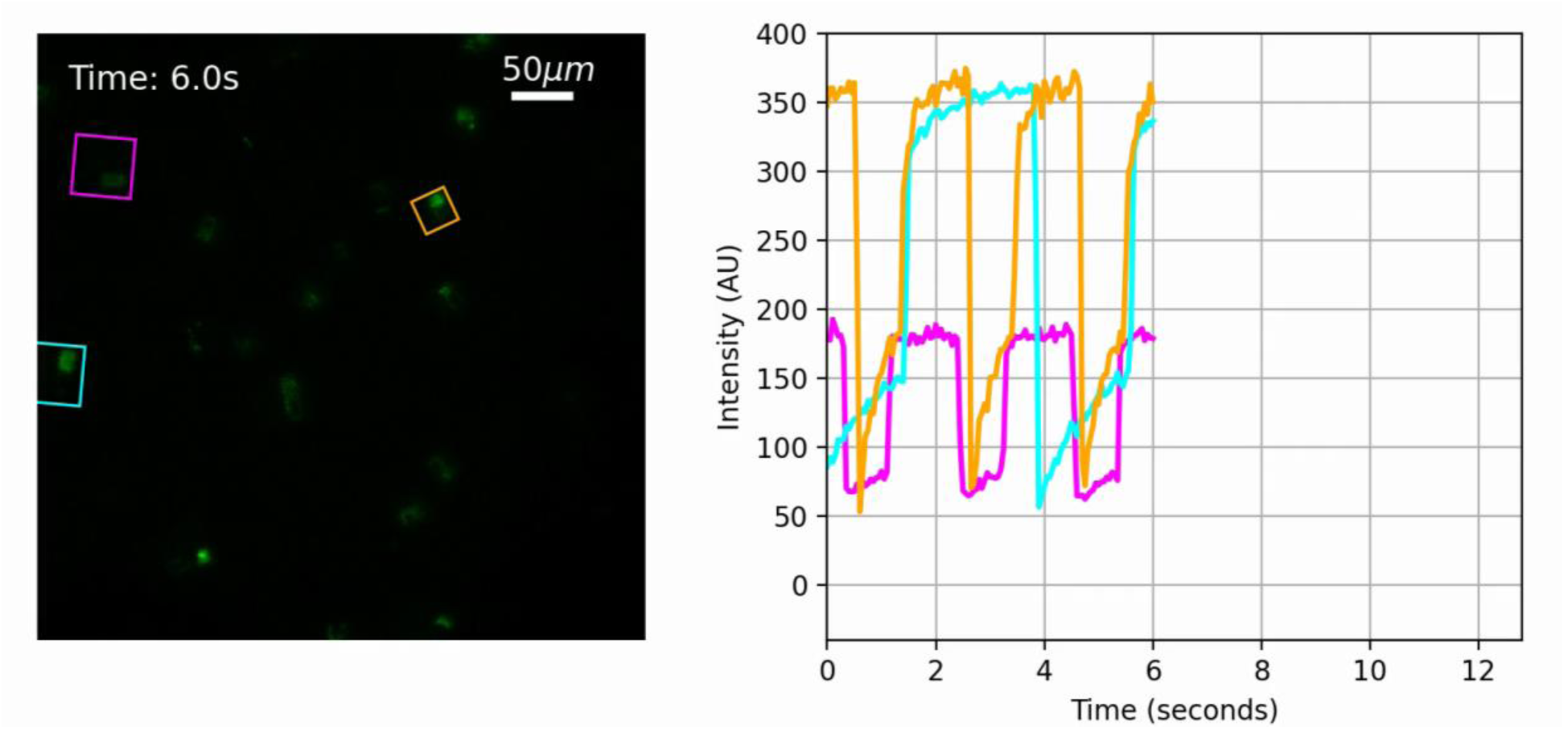
Timelapse epifluorescence microscopy of the operational motes along with a fluorescent intensity plot showing the oscillatory response spatially integrated over the mote ROI. Three motes (cyan, magenta and orange) are highlighted in the image.

